# Rapid, direct detection of bacterial Topoisomerase 1-DNA adducts by RADAR/ELISA

**DOI:** 10.1101/2020.03.09.984153

**Authors:** Devapriya Sinha, Kostantin Kiianitsa, David R. Sherman, Nancy Maizels

## Abstract

Topoisomerases are proven drug targets, but antibiotics that poison bacterial Topoisomerase 1 (Top1) have yet to be discovered. We have developed a rapid and direct assay for quantification of Top1-DNA adducts that is suitable for high throughput assays. Adducts are recovered by “RADAR fractionation”, a quick, convenient approach in which cells are lysed in chaotropic salts and detergent and nucleic acids and covalently bound adducts then precipitated with alcohol. Here we show that RADAR fractionation followed by ELISA immunodetection can quantify adducts formed by wild-type and mutant Top1 derivatives encoded by two different bacterial pathogens, *Y. pestis* and *M. tuberculosis*, expressed in *E. coli* or *M. smegmatis*, respectively. For both enzymes, quantification of adducts by RADAR/ELISA produces results comparable to the more cumbersome classical approach of CsCl density gradient fractionation. The experiments reported here establish that RADAR/ELISA assay offers a simple way to characterize Top1 mutants and analyze kinetics of adduct formation and repair. They also provide a foundation for discovery and optimization of drugs that poison bacterial Top1 using standard high-throughput approaches.

## INTRODUCTION

There is a widely recognized need to develop new drugs to treat infections, especially as many microorganisms have developed resistance to antibiotics in common use. Topoisomerases have proven to be effective drug targets not only in infectious disease but also in cancer (1,2). Topoisomerases modulate DNA topology by catalyzing cleavage and then religation of one or both strands of the duplex, forming a covalent topoisomerase-DNA complex as an obligatory reaction intermediate (3) (Fig. 1A). Accumulation of toxic Top1-DNA covalent intermediates contributes to killing of cells in which drugs or mutations impair the normal Top1 reaction cycle. Drugs that target topoisomerases cause the covalent adduct to persist, preventing release of the bound protein and religation of the DNA. This is the mechanism of cell killing by fluoroquinolone antibiotics, such as ciprofloxacin, moxifloxacin, levofloxacin, and ofloxacin, which target DNA gyrase (4); and of chemotherapeutics used to treat cancer, such as topotecan and etoposide (5,6). Mutations in topoisomerases can also impair their ability to release the covalent bond and reseal the DNA duplex.

**Figure 1.**
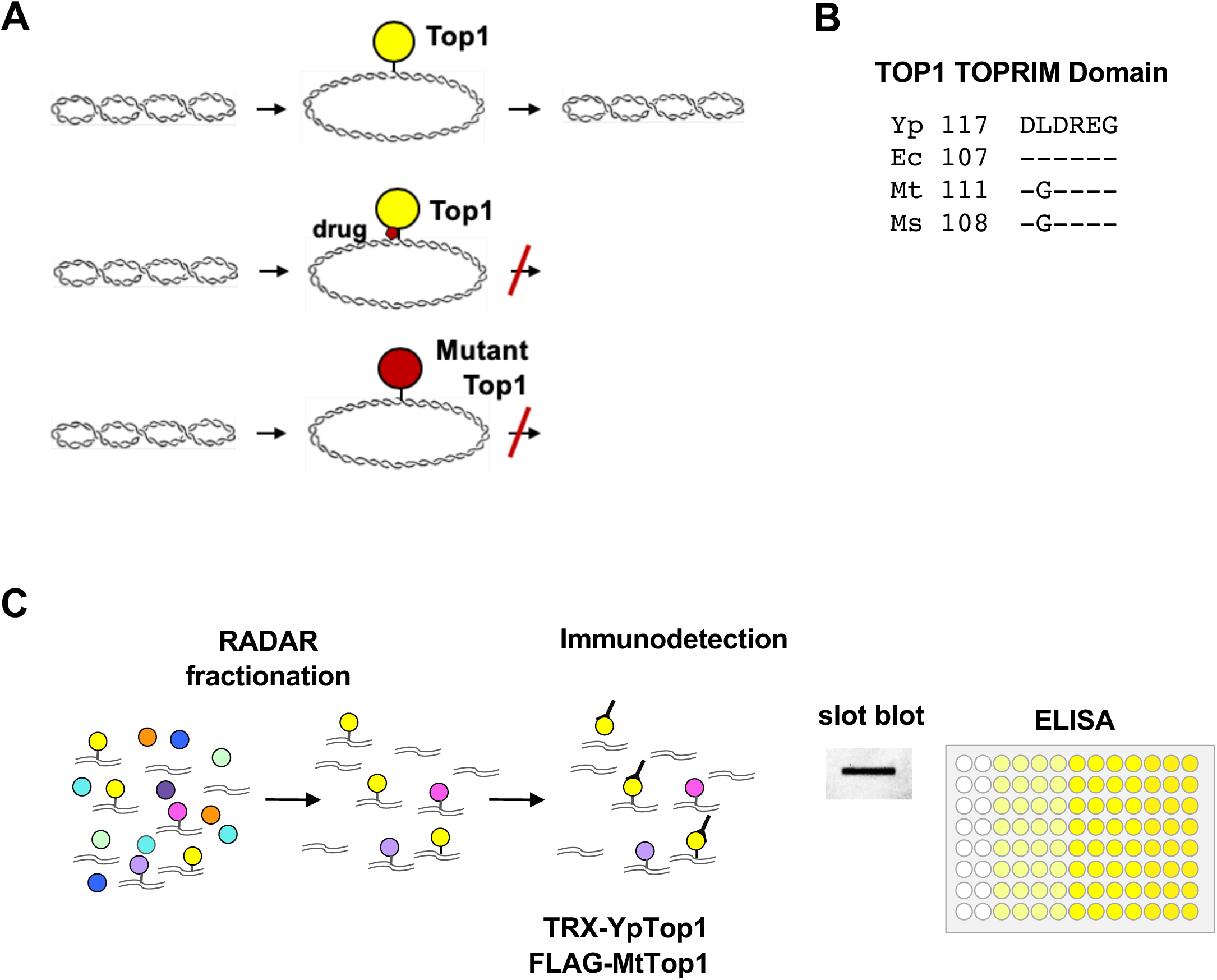
Topoisomerase 1 forms DNA adducts. (**A**) *Top line*: Bacterial topoisomerase 1 (Top1) nicks a single strand of the backbone of supercoiled DNA to form a covalent 5’-phosphoryl-tyrosine linkage and relax the supercoil. The adduct is released upon DNA religation, which may be accompanied by restored supercoiling, as shown. Drugs (*middle line*) or mutations (*bottom line*) that prevent religation (“topoisomerase poisons”) impair release of the protein-DNA adduct to cause cell killing. Images of supercoiled and relaxed circles from Wikicommons. (**B**) Conserved Top1 TOPRIM domain sequences from *Y. pestis* (Yp), *E. coli* (Ec), *M. tuberculosis* (Mt) and *M. smegmatis* (Ms). (**C**) Cell lysis in RADAR buffer (chaotropic salts and detergent) followed by alcohol precipitation enriches protein-DNA protein adducts, which are then captured for immunodetection by ELISA assay (shown) or slot blot.

The only bacterial topoisomerases targeted by drugs to date are the Type II topoisomerases, gyrase and topoisomerase IV. Efforts to identify potent small molecule inhibitors of bacterial Type IA enzymes have had only limited success thus far (7). Key to the religation step catalyzed by bacterial Top1 is coordination of a metal ion by aspartate residues of an acidic triad within the DxDxxG motif of the highly conserved Top1 TOPRIM domain (8,9) (Fig. 1B). Mutational analysis has shown that modification of the N- or C-terminal residues of this domain in Top1 from bacterial species, including *Escherichia coli, Yersinia pestis, Mycobacterium tuberculosis* and *Mycobacterium smegmatis*, will impair DNA religation, induce an SOS response and cause cell killing (10-14).

The absence of a simple, mechanism-based assay that quantifies Top1-DNA adducts formed in vivo has been a stumbling block to drug discovery and development for this target. Some features of topoisomerases themselves contribute to difficulty in systematic detection. Topoisomerase-DNA adducts are normally transient and may resolve spontaneously when drug is removed or limiting; cells may lyse in response to cell killing, releasing adducted complexes into the culture medium. The adducted enzyme may also undergo proteolytic repair, eliminating epitopes for immunodetection, as we recently demonstrated for human endogenous topoisomerase-DNA adducts by an unbiased proteomic approach (15). As the first step toward overcoming these challenges, we have developed a rapid and direct assay for quantification of topoisomerase-DNA adducts formed in living cells. The assay is based on a method referred to as “RADAR” fractionation (16,17), in which cells are lysed in chaotropic salts and detergent; nucleic acids and adducted proteins are separated from free protein by alcohol precipitation; and adducted protein is quantified by immunodetection. RADAR fractionation has enabled quantification of adducts formed by bacterial and mammalian topoisomerases (16-23) and by Pol beta (24) and other mammalian proteins (21,25). We have recently validated the ability of RADAR fractionation to enrich adduct-forming proteins from human cells by mass spectrometry (15). RADAR fractionation is cost- and time-effective, and provides a considerable improvement in throughput (100-fold or more) over the classical approach for adduct recovery by CsCl density gradient fractionation (26).

Here we describe an assay for bacterial Top1-DNA adducts that combines RADAR fractionation and ELISA assay in microplate format. We have developed this assay using as a model Top1-DNA adducts encoded by two different bacterial pathogens, *Y. pestis* and *M. tuberculosis*, expressing epitope-tagged Top1 from inducible constructs. This enables detection by highly specific commercially available antibodies, circumventing the need to identify and characterize antibodies specific to each target of interest. We apply the RADAR/ELISA assay to quantify accumulation of DNA adducts formed by the highly toxic *Y. pestis* mutant, YpTop1-D117N, expressed in *E. coli* cells. The Z’ factor of the assay is >0.5, suitable for high throughput applications. We further demonstrate that RADAR lysis disrupts the normally challenging mycobacteria cell wall and enables recovery and quantification of adducts formed by MtTop1-DNA expressed in *M. smegmatis*. These experiments provide a foundation for discovery and optimization of drugs that poison bacterial Top1 using standard high-throughput approaches.

## MATERIALS AND METHODS

### Cloning and site-directed mutagenesis of MtTop1 constructs

Gateway LR clonase (Thermo-Fisher) was used to clone the MtTop1 coding sequence from the pENTR vector (pENTR:3646c; (27)) into the pDTNF expression vector (pDTNF:3646c), fusing a FLAG tag to the N-terminal of Top1. The D111A, D111N, and G116S mutants of MtTop1 were generated by QuikChange using *Pfu* Turbo (Thermo-Fisher).

### Cell culture, Top1 induction, and viability assays

*E. coli* strain MG1655 with chromosomally integrated YpTop1 WT (BWYTOP) or YpTop1-D117N (BW117N) bearing an N-terminal TRX tag was a gift from Prof Yuk-Ching Tse-Dinh (Florida International University, USA). Cells were cultured at 37°C in LB medium containing 25 μg/ml chloramphenicol. Top1 expression was induced in exponentially growing cultures (OD_600_=0.3) by addition of arabinose (Sigma Aldrich). Cell division was monitored by measuring OD_600_.

*M. smegmatis* (mc^2^155) strain was cultured in 7H9 medium with ADC supplement (HiMedia) as described (28). *M. smegmatis* was transformed using a MicroPulser™ Electroporator Bio-Rad), and transformed cells selected and propagated in medium containing 200 μg/ml hygromycin B (Thermo-Fisher). MtTop1 expression was induced in exponentially growing (OD_600_=0.5) cultures of *M. smegmatis* by addition of 50 ng/ml anhydrotetracycline (ATc; Takara-Clontech) (29). Cell survival was quantified by culturing cells on LB plates containing 200 μg/ml hygromycin B for 3 days at 37°C.

*M. tuberculosis* H37Rv (ATCC 25618) strain was grown in 7H9 medium with ADC supplement (HiMedia) as described (28). Up to 2×10^9^ Mtb bacilli were incubated in RADAR buffer for 15 min at 65°C with occasional vortexing, washed once in water, plated, and then incubated for four weeks at 37°C.

### RADAR lysis reagent for bacteria

To perform lysis and RADAR fractionation of bacteria, we used a reagent developed for proteomic analyses of nucleoprotein adducts in human cells (26), with some modifications. The lysis solution for bacteria (LSB) consisted of 5 M guanidinium isothiocyanate (GTC), 1% Sarkosyl, 1% 2-mercaptoethanol, 20 mM EDTA, 20 mM Tris-HCl (pH 8.0) and 0.1 M sodium acetate (pH 5.3), adjusted to final pH 6.5 with NaOH. The solution was filtered through 0.1 µm PES membrane (VWR) and stored in the dark at room temperature. 2-mercaptoethanol was freshly added before use. For alkaline lysis and fractionation, LSB was supplemented with 5 N NaOH to the desired concentration.

### RADAR/slot blots of Top1-DNA adducts in *E. coli*

RADAR fractionation was carried out as previously described (16,17) with some modifications. *E. coli* cells (OD_600_ =0.3-0.5) bearing YpTop1 expression plasmids were cultured in 20 ml of LB broth with or without arabinose inducer and harvested by centrifugation at 3,500 rpm for 10 min. The pellet (approximately 2×10^10^ cells) was lysed in 500 μl LSB supplemented with 0.25 M NaOH, and incubated at 60°C for 15 min. Samples were then sonicated with 30s pulses, 100 amplitude, for 3-4 cycles, and the extract clarified by centrifugation at 21,000 g for 10 min. To 450 µl of resulting supernatant were added 150 µl of 8 M LiCl (final concentration 2 M) and 600 µl isopropanol (equal volume), followed by centrifugation at 21,000 g for 10 min. The resulting pellet was washed thrice with 75% ethanol, briefly air dried, resuspended in 400 µl freshly prepared 8 mM NaOH, dissolved on the Thermomixer at 2000 rpm at room temperature and neutralized by addition of 1 M HEPES free acid solution (8 µl). To reduce background of non-covalently bound overexpressed Top1, samples (100 µl) were treated with RNase A for 30 min at 37°C (final concentration 20 µg/ml), supplemented with 350 µl LS1 and 150 µl of 8M LiCl solution (final concentration 2M LiCl), reprecipitated with equal volume of isopropanol and resuspended in 8 mM NaOH. DNA concentrations were measured with a Qubit assay. Typical recovery was 30-40 μg of DNA from 1.8×10^10^ *E. coli* cells. Prior to slot blotting, 100 µl of sample was removed and digested with 0.5 μl (12.5 units) Benzonase (Novagen) in the presence of 2 mM MgCl_2_ for 30 min at 37°C.

For slot blotting, 100 μl of sample containing 0.5-1 µg of Benzonase-digested DNA in 25 mM sodium phosphate buffer (pH 6.5) was applied to a nitrocellulose membrane (Bio-Rad) pre-wet in the same buffer using a vacuum slot blot manifold (Bio-Rad). Membranes were blocked in 0.5% alkali soluble casein (Novagen) in 10 ml TBST (10 mM Tris-EDTA pH 7.5, 0.1 M NaCl, 0.05% Tween 20) for 1 hr. TRX-tagged YpTop1 was detected with polyclonal rabbit anti-thioredoxin antibodies (Abcam, ab26320, 1:1,000). Membranes were incubated with primary antibodies for 3 hr, washed in TBST and incubated for 1hr with HRP-conjugated anti-rabbit secondary antibodies (Thermo-Fisher; 1:10,000). All antibodies were diluted in the blocking solution, and all incubations were performed at room temperature. Membranes were developed using Super Signal West Dura (Thermo-Fisher) and imaged on a Bio-Rad Chemidoc XRS Plus Analyzer.

### CsCl density gradient fractionation of YpTop1-DNA adducts in *E. coli*

CsCl density gradient fractionation and adduct quantification were based on a protocol developed for bacterial gyrase and topoisomerase IV adducts (30), with some modifications. Briefly, a 50 ml culture (5×10^10^ cells) was pelleted, resuspended in 3 ml buffer containing TE (10 mM Tris-HCl pH 7.5, 1 mM EDTA pH 8.0), 1x protease inhibitor cocktail (Thermo Scientific), and lysozyme (Sigma Aldrich) added to a final concentration of 0.1 mg/ml. Samples were incubated on ice for 10 min, sarkosyl added to final concentration 1%, then following an additional 30 min incubation on ice DNA was sheared by expulsion through 22G1/2 needles followed by centrifugation at 21,000 g for 10 min. Supernatants (3 ml) were loaded on the top of step gradients preformed in polyallomer tubes (14 by 89 mm, Beckman), containing 2 ml each of 1.82, 1.72, 1.50, and 1.37 g/ml CsCl (Sigma Aldrich) in 10 mM Tris, 1 mM EDTA, pH 8.0. Following centrifugation in a Beckman SW41Ti rotor at 31,000 rpm for 20 hr at 20°C, the bottom of the tube was punctured, and 14 fractions were collected. DNA was quantified by Qubit assay. To compare levels of adducts formed by WT and mutant YpTop1, equal amounts of DNA from each fraction (1,000 ng) were prepared to load onto the membrane; prior to slot blotting, samples were digested with 0.25 μl Benzonase in the presence of 2 mM MgCl_2_ for 30 min at 37°C. TRX-tagged YpTop1 was detected, as described for YpTop1 RADAR/slot blots.

### RADAR/ELISA assays of YpTop1-DNA adducts in *E. coli*

For RADAR fractionation in 96-well microtiter plates, cells (4×10^7^) were cultured in 50 µl medium and protein expression induced with 0.02% arabinose. Cells were pelleted by centrifugation for 5 min at 2,700 g, resuspended in 50 μl 1x FastBreak™ Cell Lysis Reagent (Promega) supplemented with RNase A (20 µg/ml) and incubated at 37°C for 5 min, followed by addition of 50 μl LSB premixed with 12 M LiCl (1:6, final concentration 1 M LiCl). After addition of 100 μl isopropanol (equal volume) DNA was precipitated by centrifugation for 10 min at 2,700 g. Pellets were washed with 200 µl 75% ethanol, briefly air dried and solubilized on the Thermomixer (20 min, 2,000 rpm) in 25 μl 8 mM NaOH, followed by neutralization with 1 M HEPES. Typical recovery following culture and fractionation in 96 well plates was 500-700 ng of DNA from 10^8^ *E. coli* cells.

For the homogeneous assay, cells were not pelleted. Instead, 5 μl 10x FastBreak™ Cell Lysis Reagent supplemented with RNase A was added directly to 50 µl of bacterial culture and incubated on the Thermomixer for 5 min at 37°C, then 50 µl LSB-LiCl mixture was added and all subsequent steps carried out as above.

Sandwich ELISAs in 96-well format were used to quantify TRX-tagged YpTop1 adducts in samples containing the equivalent of 100 ng DNA per well. Prior to ELISA, DNA was digested with Benzonase in the presence of 2mM MgCl_2_. Samples were applied to ELISA plates (Nunc Poly-sorb) pre-coated with rabbit polyclonal anti-TRX capture antibodies (EpiGentek #A57734, 0.5 μg/ml). Primary detection of TRX-tagged YpTop1 was with primary murine monoclonal anti-TRX (BioLegend #658902, 1μg/ml),. Secondary detection used biotin-conjugated anti-mouse IgG (BioLegend *#*405303, 0.5 μg/ml), followed by HRP-Streptavidin conjugates (BioLegend #405210, 1:1,000). All antibody dilutions were in 1x ELISA assay diluent (BioLegend).

Z’ factors were calculated for assays carried out at different times post-induction (0-5.5 hr) to test the effectiveness of assay conditions and to determine the optimum time at which to assay adducts. Z’ was calculated using the formula: Z’= 1-(3σ_p_ + 3σ_b_) (μ_b_-μ_p_), where σ_p_ and σ_b_ are the standard deviation of the signals of YpTop1 D117N and YpTop1 WT, respectively; μ_b_ represents the mean of the signal obtained from YpTop1 WT and μ_p_ is the mean of YpTop1-D117N.

### RADAR/ELISA assays of MtTop1-DNA adducts in *M. smegmatis*

After induction of MtTop1 expression with anhydrotetracycline (ATc, Sigma; 50 ng/ml), 2 ml cultures (approximately 2.5×10^9^ bacilli) were collected and cells harvested by centrifugation at 21,000 g for 2 min. Pellets were washed with 1 ml sterile H_2_O, centrifugated as above, and resuspended in 100 µl TE. Suspensions were then sonicated for 3-4 cycles of 30 sec pulses, 100 amplitude, and treated with RNase A (10 µg/ml) for 30 min at 37°C. Then, 400 µl LSB was added, followed by 15 min incubation on the Thermomixer at 60°C. Extracts were clarified by centrifugation at 21,000 g for 10 min, supernatants (450 µl) transferred to new tubes and supplemented with LiCl (final concentration 2M LiCl). After mixing with equal volume of isopropanol, DNA was precipitated at 21,000 g for 10 min. DNA pellets were washed with 75% ethanol, briefly dried in air, and 50 μl of 8 mM NaOH was added prior to solubilization on a Thermomixer followed by neutralization with 1 μl 1M HEPES. To perform fractionation in 96 deep-well plate format, aliquots of 0.5-2.5×10^9^ *M. smegmatis* cells were processed as above, except all centrifugations were performed at 2,700 g. DNA concentrations were determined by Qubit assay. Prior to ELISA, DNA was digested with Benzonase in the presence of 2 mM MgCl_2._

For direct ELISA of FLAG-tagged MtTop1 adducts isolated from *M. smegmatis*, samples were adjusted to 1xELISA coating buffer (BioLegend) and applied to untreated ELISA plates. Samples were absorbed for 2 hr at room temperature or overnight at 4°C. Wells were washed 4 times with 100 μl of 1xPBS, 5 min per wash, then blocked with 100 μl of 1xELISA assay diluent (BioLegend). Following incubation with 1 μg/ml mouse monoclonal anti-DDK (FLAG) antibody (60 µl per well, 3 hr at room temperature), plates were washed 4 times for 5 min, then incubated for 45 min with 60 µl HRP-conjugated secondary goat anti-mouse IgG (Thermo Scientific, 1:5,000) and washed as above. Signal detection was performed using TMB High Sensitivity Solution (BioLegend, # 421501) according to manufacturer’s instructions. Absorbance was read at 450 and 570 nm, and the A570 reading was subtracted as background from the A450 signal to correct for background.

### CsCl density gradient fractionation of MtTop1-DNA adducts in *M. smegmatis*

Bacteria (ca. 5×10^10^ cells) were harvested by centrifugation, washed in water and resuspended in 3 ml TE with 1x protease inhibitor cocktail. Samples were sonicated on ice for 4 cycles of 30 Amp and 30 sec pulse time on/off, followed by addition of RNase A (Thermo Fisher) to a final concentration of 10 μg/ml and sarkosyl to a final concentration of 1%, after which samples were incubated for 30 min on ice.

Density gradient centrifugation, fraction collection and DNA quantification were performed as described for YpTop1 adducts in *E.coli*. Prior to dot blotting, 10 µl of each fraction was digested with 0.25 μl Benzonase in the presence of 2 mM MgCl_2_ for 30 min at 37°C. FLAG-tagged MtTop1 was detected with anti-DDK (FLAG) antibodies (Origene, TA50011, 1:2,500). Membranes were incubated with primary antibodies at room temperature for 3 hr, washed in TBST and incubated for 1 hr with HRP-conjugated anti-mouse secondary antibodies (Thermo-Fisher; 1:10,000). Signal was detected, as described for the RADAR/slot blots of YpTop1-DNA adducts.

## RESULTS

### Adducts of cytotoxic YpTop1-D117N accumulate upon expression in *E. coli*

We assayed adducts formed by YpTop1 WT and YpTop1-D117N bearing N-terminal thioredoxin (TRX) tags, expressed in *E. coli.* YpTop1-D117N bears a mutation in the TOPRIM domain (Fig. 1B) that eliminates a negatively charged residue required for Mg^2+^ interaction, rendering the protein defective in DNA religation and extremely cytotoxic upon expression in *E. coli* cells (11,12). YpTop1 WT and YpTop1-D117N were expressed from an arabinose-inducible promoter, and OD_600_ of the cultures determined during 4 hr following induction of protein expression by addition of arabinose. Induction did not affect viability of cells expressing YpTop1 WT, but a clear drop in OD_600_ occurred in cells expressing YpTop1-D117N (Fig. 2A). Plating assays carried out at 3 hr postinduction confirmed that expression of YpTop1-D117N, but not YpTop1 WT diminished cell viability (Fig. 2B). Analysis of DNA adducts by RADAR/slot blot identified only a faint signal in samples from uninduced cultures or from induced cultures expressing YpTop1 WT, but an intense signal in samples from induced cultures expressing YpTop1-D117N (Fig. 3C). Thus, impaired survival caused by induction of YpTop-D117N expression in *E. coli* correlated with accumulation of DNA adducts as assayed by RADAR/slot blot.

**Figure 2.**
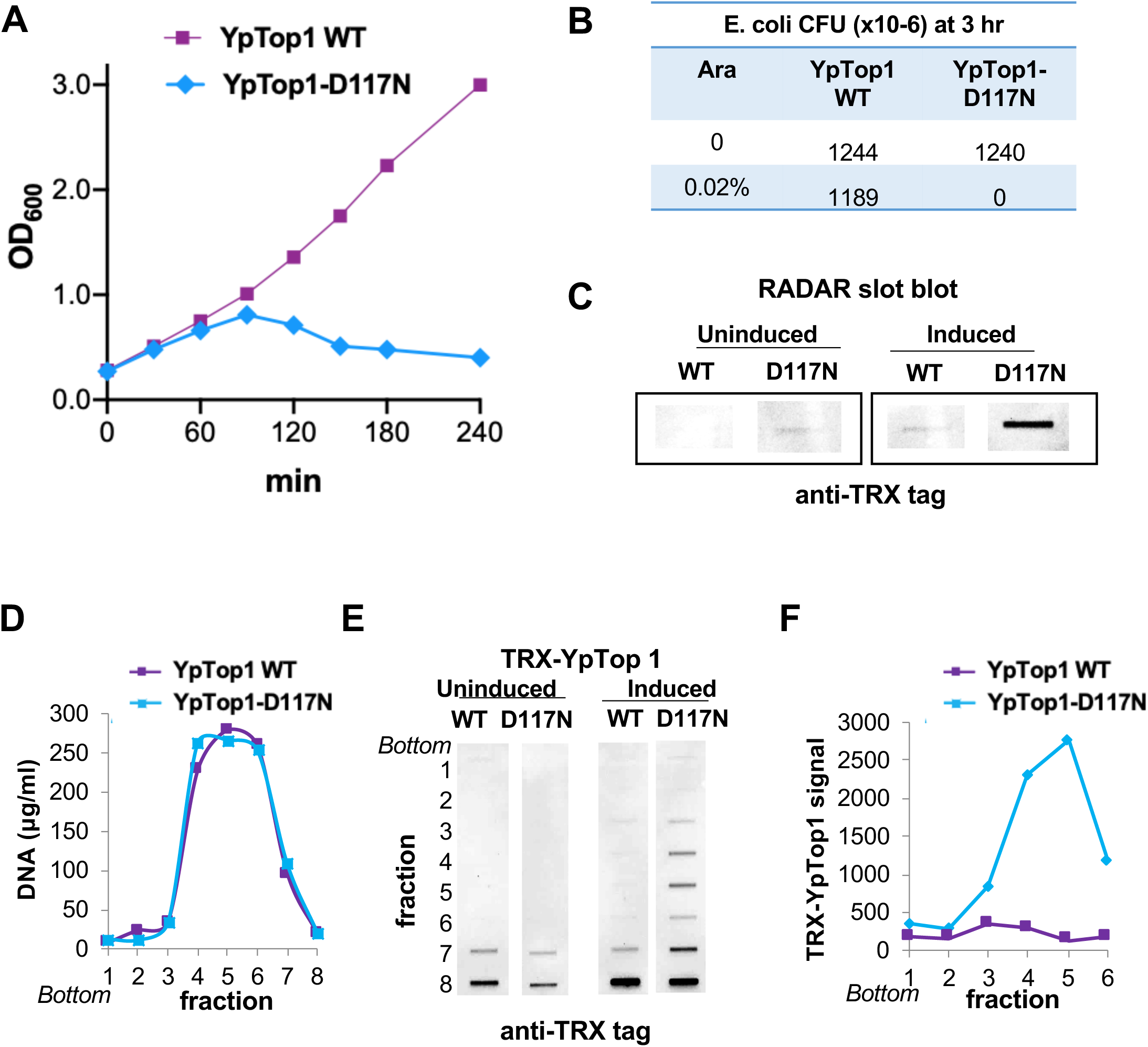
Quantification of YpTop1 WT and YpTop1-D117N DNA adducts by RADAR slot blot and CsCl buoyant density fractionation. (**A**) Growth kinetics (OD_600_) of cultures of *E. coli* bearing arabinose-inducible expression clones for TRX-tagged YpTop1 WT and YpTop1-D117N, assayed from 0-240 minutes after induction with indicated concentration of arabinose. (**B**) Representative assay of CFU recovered from *E. coli* expressing YpTop1 WT and YpTop1-D117N, assayed at 180 minutes of culture with 0.02% arabinose. (**C**) Slot blot of RADAR-fractionated extracts isolated from *E. coli* at 150 min post-induction of expression of YpTop1 WT or YpTop1-D117N by culture with arabinose. Immunodetection was performed with antibodies to TRX epitope. (**D**) Quantification of DNA recovery by CsCl gradient fractionation of extracts of cells that were uninduced or induced by 150 min culture with arabinose. (**E**) Slot blot of extracts of uninduced and induced cells fractionated by CsCl density gradient centrifugation. *Bottom*, bottom of gradient. (**F**) Quantification of TRX-tagged YpTop1 derivatives recovered by CsCl gradient fractionation and detected by slot blot.

**Figure 3.**
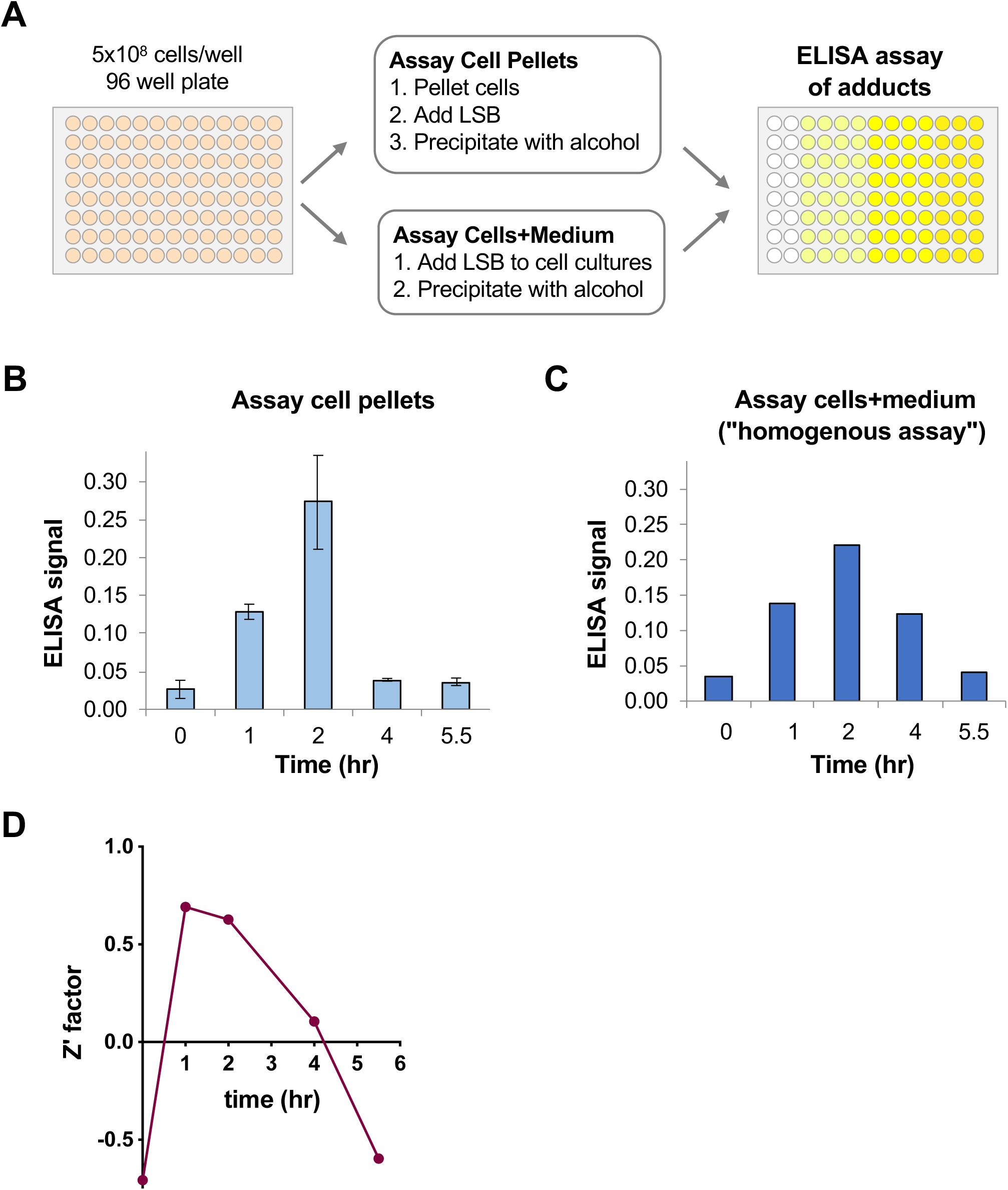
Reproducible RADAR/ELISA assay in microtiter format. (**A**) Flowchart for adduct recovery from cells cultured in microtiter format. (**B**) RADAR/ELISA assay of YpTop1 signal from pelleted cells expressing YpTop1-D117N at indicated times after induction of expression with arabinose (0.2%). (**C**) Homogeneous RADAR/ELISA assay of YpTop1 signal from entire culture (cells and media) expressing YpTop1-D117N at indicated times after induction of expression with arabinose (0.2%). (**D** Z’ factors of homogenous Top1 RADAR/ELISA assays calculated at indicated times post-induction of protein expression.

We used the classical approach of CsCl density gradient fractionation (26) to confirm the results of RADAR/slot blot. Extracts of cultures expressing TRX-tagged YpTop1 WT or YpTop1-D117N, uninduced or induced by 150 min culture with arabinose, were fractionated on a CsCl step gradient, and the DNA concentration of each fraction was measured and plotted, showing that DNA peaked in fractions 4-6 (Fig. 2D). Slot blots showed that signals from free protein were evident in fractions 7-8 of all samples, which are near the top of the gradient (Fig. 2E). The free protein signal increased in response to induction of Top1 expression with arabinose, as expected. The DNA peak-containing fractions of intermediate density (fractions 4-6) exhibited clear signals only in arabinose-induced cultures, with a stronger signal evident in the fractions from cells expressing YpTop1-D117N (Fig. 2E, right). To assess adduct recovery from induced cells, the Top1 signal was quantified by densitometry (Fig. 2F). This showed that more adducts accumulated in *E. coli* cells expressing YpTop1-D117N mutant than YpTop1 WT, validating the results of the RADAR/slot blot.

### Reproducible RADAR/ELISA assay in microtiter format

To further streamline RADAR quantification of adducts, we scaled down cell numbers to enable adduct quantification in microtiter plates by RADAR/ELISA assay (17). As outlined in Fig. 3A, 5×10^8^ cells were cultured, Top1 expression induced with arabinose, pelleted, lysed by treatment with chaotropic salts and detergent, and the sample then alcohol precipitated to recover nucleic acids and covalently bound proteins, resuspended, and finally aliquoted to quantify DNA recovery by Qubit assay and detect the Top1 signal by ELISA. The microtiter plate format allows multiple samples to be processed in parallel, facilitating kinetic analysis of adduct accumulation. RADAR/ELISA assays of adducts during the first 5.5 hr after arabinose induction of YpTop1 expression showed that the signal increased through 2 hr and dropped dramatically at 4 hr (Fig. 3B). This timing correlated with the reduction of OD_600_ in the culture expressing YpTop1-D117N (Fig. 2B), suggesting that adducts might be released into the medium upon cell lysis.

To assay adducts in both intact cells and the culture medium, we devised a “homogenous assay” in which cells were not pelleted prior to addition of lysis buffer. Instead, cell lysis and ethanol precipitation were carried out on the entire contents of each well of a 96-well micro-titer plate. RADAR/ELISA assay showed that the protein signal as detected by this approach peaked at 2 hr and then decreased gradually (Fig. 3C), evidence of reduced sensitivity of the assay to cell lysis.

To establish whether the homogeneous RADAR/ELISA assay was sufficiently quantitative and reproducible for high throughput applications, we determined Z’ factors for different time points of Top1 induction. Assays with Z’>0.5 are considered excellent, and this criterion was satisfied by homogeneous RADAR/ELISA assay of Top1-DNA adducts at 1-2 hr post-induction (Fig. 3D).

### RADAR lysis conditions efficiently recover *DNA from Mycobacterium* sp

To extend the RADAR/ELISA assay to a context other than *E. coli*, we focused on *Mycobacteria. Mycobacterium tuberculosis* (Mtb) causes tuberculosis (TB), a major challenge to human health worldwide and the deadliest infectious disease, after AIDS (WHO, 2019; https://www.who.int/tb/en/). Mtb encodes a single type IA DNA topoisomerase (MtTop1) that is crucial for viability and predicted to be a drug target (31-33). Mycobacteria are notoriously difficult to disrupt, as the cells are protected by a tough outer layer composed of lipids, mycolic acids, polysaccharides (arabinoglycan) and peptidoglycans that make them highly resistant to lysis by standard chemical or enzymatic approaches. Strikingly, treatment with LSB at 65°C was toxic to the virulent Mtb H37Rv (ATCC 25618) strain, which would enable treated Mtb cells to be handled outside a BSL-3 facility. Mtb are very slow-growing, so for assay development we turned instead to *M. smegmatis* and BCG. We found that we were conveniently able to achieve DNA yields on the order of 20% from both bacteria by incubation of cells at 65°C for 10 min in RADAR LSB, followed by isopropanol precipitation and centrifugation at 21,000 g for 10 min using a standard benchtop microfuge (Fig. 4A). DNA yields from *M. smegmatis* or BCG were considerably improved by sonication, which is useful for large-scale preparation but requires specialized equipment for analysis of small volumes in microplate format.

**Figure 4.**
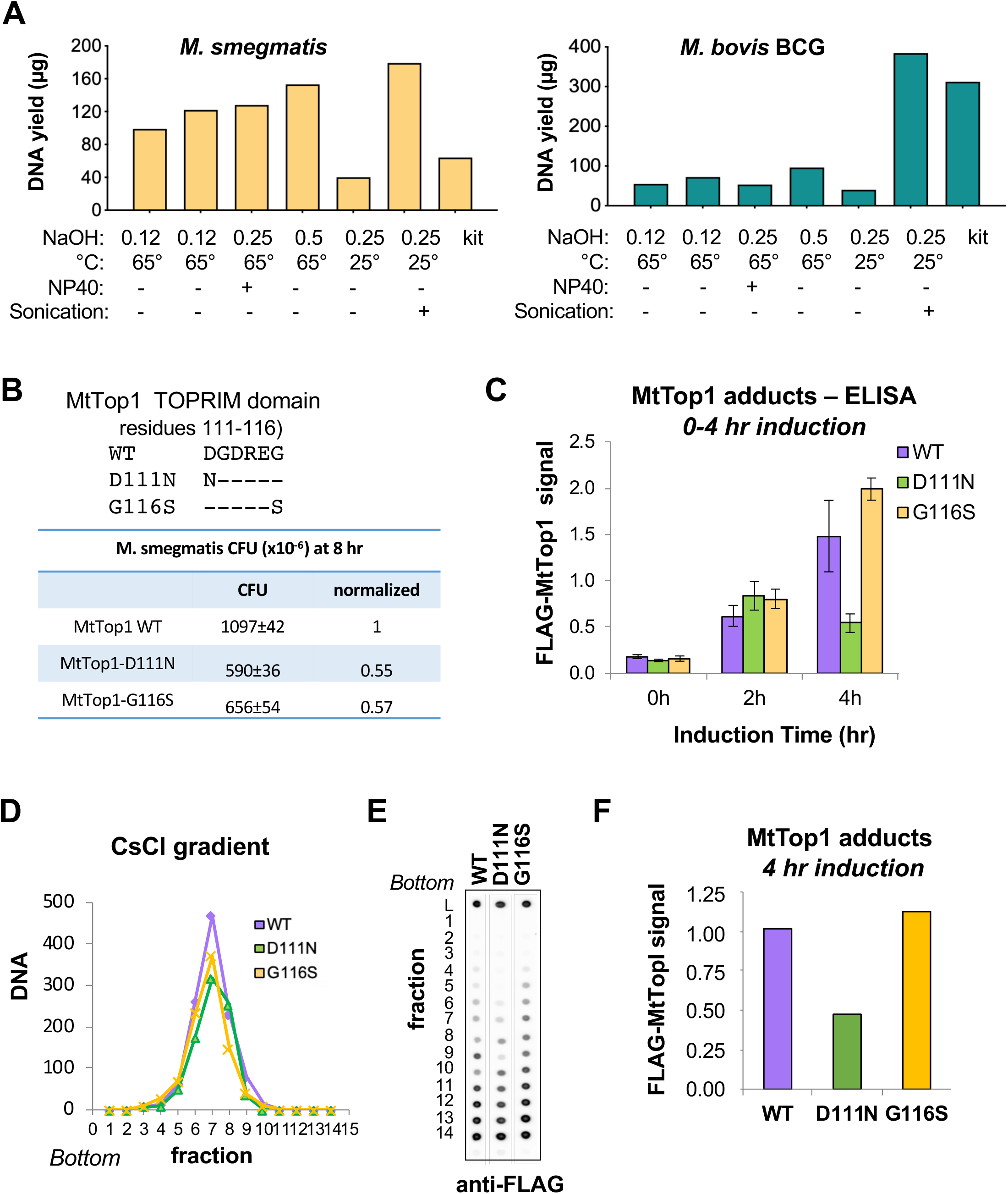
RADAR/ELISA quantifies MtTop1 DNA adducts in *M. smegmatis*. (**A**) Recovery of DNA from (*left*) *M. smegmatis* (ng/10^8^ cells) or (*right*) BCG. Bacterial lysis was performed in LSB supplemented with NaOH at indicated concentrations and in the absence/presence 2% NP-40, followed by alcohol precipitation to recover DNA. Kit: DNA was isolated using the commercial MagJET Genomic DNA Kit. (**B**) *Above*, TOPRIM domain sequences of WT and mutant MtTop1. *Below*, CFU recovered from *M. smegmatis* expressing MtTop1 WT and indicated mutants at 8 hr after induction of MtTop1 expression with ATc. Colonies were counted after 3 days growth at 37°C on plates containing 200 µg/ml hygromycin B. Shown are absolute values and survival normalized to cells expressing MtTop1 WT. (**C**) RADAR/ELISA assay of FLAG-MtTop1 signal from pelleted cells expressing WT and mutant FLAG-MtTop1 at indicated times after induction of expression with ATc (50 ng/ml). (**D**) Quantification of DNA recovered in CsCl gradient fractions of extracts of *M. smegmatis* cells expressing FLAG-MtTop1 WT and indicated mutants following 4 hr induction with ATc. *Bottom*, bottom of gradient. (**E**) Slot blot detection of FLAG-tagged MtTop1 derivatives in indicated CsCl density gradient fractions. *Bottom*, bottom of gradient. (**F**) Quantification of recovery of FLAG-MtTop1 from WT and mutant derivatives. Each signal represents the total from fractions 5-9, which contain the peak of DNA.

### RADAR/ELISA detects MtTop1 DNA adducts in *M. smegmatis* cells

To analyze adduct formation by MtTop1 expressed in *M. smegmatis*, we modified a construct for tetracycline-inducible expression of MtTop1 to carry an N-terminal FLAG (DDK) tag for immunodetection. We then generated mutations in MtTop1 at two positions reported to cause toxicity in other bacterial Top1 proteins (Fig. 4B, above). These included a D111N mutation, corresponding to the YpTop1-D117N mutation toxic in *E. coli* (11); and a G116S mutation, corresponding to mutations in Yp or EcTop1 that caused a dramatic decrease in cell viability and induced an SOS response in *E. coli* (10). Cloned DNA was introduced into *M. smegmatis* (mc^2^155), cells were cultured to early log phase (OD_600_∼0.5), and MtTop1 expression induced by addition of anhydro-tetracycline (ATc, 50 ng/ml). Analysis of viable CFU in cells plated at 8 hr post-induction found that expression of MtTop1-D111N or MtTop1-G116S reduced viability only 2-fold, as shown by analysis of CFU recovered in two independent experiments (Fig. 4B, below). The limited toxicity may reflect structural differences between MtTop1 and EcTop1 (9) and/or physiological differences between *E. coli* and *M. smegmatis*.

To quantify adducts by RADAR/ELISA assay, cells expressing MtTop1 WT, D111N or G116S were cultured to OD=0.5, then ATc added and cells cultured 0, 2 or 4 hr post-induction. Adducts were then enriched by RADAR fractionation, and MtTop1-DNA complexes detected by direct ELISA with antibody to the N-terminal FLAG tag (Fig. 4C). Comparable ELISA signals were evident in RADAR fractions of cells expressing WT and mutant enzymes prior to induction of MtTop1 expression (t=0). The signal from extracts of cells expressing MtTop1 WT or the G116S mutant increased through the 4 hr induction period, and the signal from extracts of cells expressing the D111N mutant increased at 2 hr then dropped slightly by 4 hr.

We carried out CsCl buoyant density fractionation to confirm results of RADAR/ELISA. *M. smegmatis* cells expressing WT or mutant MtTop1 were cultured for 4 hr with 50 ng/ml ATc, pelleted, lysed by sonication, and samples resolved on CsCl step gradients. Fractions were collected by puncturing the bottom of each tube. Fractions 5-9 were shown to contain the peak of DNA in each sample (Fig. 4D). Proteins in aliquots of each fraction were captured on a slot blot and FLAG-tagged Top1 detected with anti-FLAG tag antibodies (Fig. 4E). Adduct recovery was quantified by summing signals determined by densitometry of DNA-containing fractions (5-9) and normalizing to DNA concentration (Fig. 4F). FLAG-Top1 signals at 4 hr post-induction determined by density gradient centrifugation were found to be comparable to those determined by RADAR/ELISA.

## DISCUSSION

Here we report a straightforward assay that directly quantifies bacterial Top1-DNA adducts by ELISA assay of RADAR-fractionated cells. We demonstrate that RADAR/ELISA assay can be used to quantify adducts formed by Top1 encoded by two different pathogenic bacteria, *Y. pestis* and Mtb.

Analysis of YpTop1 expressed in E. coli showed that RADAR/ELISA assay offers a simple approach for characterization of Top1 mutants YpTop1 WT and its highly toxic mutant derivative YpTop1-D117N were expressed in *E. coli*. Adducts formed by the YpTop1-D117N mutant were detected at greater levels than YpTop1 WT adducts, as assayed by RADAR/slot blot, RADAR/ELISA, or CsCl fractionation. The increased level of adducts suggests that this mutant, which is deficient in religation, mimics the effect of treatment with topoisomerase poisons.

The results presented here report analysis of N-terminal tagged recombinant Top1 expressed from inducible promoters and detected with epitope-specific tags. The tag enables assays of adducts formed by topoisomerases for which suitable, specific antibodies are not available commercially. The essay is optimized for bacterial Top1, but it can be extended to other epitope-tagged topoisomerase targets, such as gyrases. The utility of a tag will be limited if proteolytic cleavage occurs at sites between the tag and the Top1-DNA covalent bond. We have recently shown that proteolytic repair of human topoisomerase 1 can be assessed by RADAR fractionation combined with mass spectrometry (15). That approach could be readily adopted to determine whether bacterial topoisomerase-DNA adducts were targets of proteolytic cleavage.

Cell lysis may accompany cell killing that results from drug treatment. We found that fractionation of adducts from both cells and culture medium offers a work-around, although even with this approach adduct detection diminished at later times, likely reflecting proteolysis. This highlights the importance of kinetic analysis in optimizing assay conditions. RADAR/ELISA simplifies kinetic analyses by enabling large numbers of samples to be assayed in parallel.

RADAR/ELISA proved able to detect Top1-DNA adducts fractionated from either *E. coli* or *M. smegmatis* cells, providing a foundation for extension of the assay to other bacteria. The tough cell wall of Mycobacteria presents a considerable challenge to many biochemical approaches, but sufficient DNA was recovered by RADAR fractionation to enable quantification of adducts. This suggests that the approach can be extended to other bacteria. The results reported here thus provide a general foundation for discovery and optimization of drugs that poison bacterial Top1 using standard high-throughput approaches.

## SUPPLEMENTARY DATA

none.

## ACKNOWLEDGEMENTS

We thank Dr. Yuk-Ching Tse-Dinh for providing *E. coli* strains bearing YpTop1 and for valuable advice on experimental design.

